# Hsp90 and its co-chaperone Sti1 control TDP-43 misfolding and toxicity

**DOI:** 10.1101/2020.10.08.331173

**Authors:** Lilian Tsai-Wei Lin, Abdul Razzaq, Sonja E. Di Gregorio, Soojie Hong, Brendan Charles, Marilene H. Lopes, Flavio Beraldo, Vania F. Prado, Marco A.M. Prado, Martin L. Duennwald

**Affiliations:** Department of Pathology and Laboratory Medicine, Schulich School of Medicine and Dentistry, University of Western Ontario, Canada; Robarts Research Institute, Schulich School of Medicine and Dentistry, University of Western Ontario, Canada; Department of Anatomy & Cell Biology, Schulich School of Medicine and Dentistry, University of Western Ontario, Canada; Department of Physiology and Pharmacology, Schulich School of Medicine and Dentistry, University of Western Ontario, Canada; Department of Cell and Developmental Biology, Institute of Biomedical Sciences, University of São Paulo, São Paulo, Brazil

## Abstract

Protein misfolding is a central feature of most neurodegenerative diseases. Molecular chaperones can modulate the toxicity associated with protein misfolding, but it remains elusive which molecular chaperones and co-chaperones interact with specific misfolded proteins. TDP-43 misfolding and inclusion formation is a hallmark of amyotrophic lateral sclerosis (ALS) and other neurodegenerative diseases. Using yeast and mammalian neuronal cells we find that Hsp90 and its co-chaperones have a strong capacity to alter TDP-43 misfolding, inclusion formation, aggregation, and cellular toxicity. Our data also demonstrate that impaired Hsp90 function sensitizes cells to TDP-43 toxicity. We further show that the co-chaperone Sti1 specifically interacts with and modulates TDP-43 toxicity in a dose-dependent manner. Our study thus uncovers a previously unrecognized tie between Hsp90, Sti1, TDP-43 misfolding, and its cellular toxicity.

## Introduction

Protein misfolding and aggregation are hallmarks of protein conformational diseases, including neurodegenerative diseases, such as Alzheimer’s disease (AD), Parkinson’s disease (PD), Huntington’s disease (HD), Frontal Temporal Dementia (FTD), and amyotrophic lateral sclerosis (ALS) (1–3). ALS is characterized by death of upper and lower motor neurons causing progressive loss of muscle function, eventually leading to fatal paralysis of the respiratory system (4). In the majority of ALS cases (∼97%), the transactive response element <u>D</u>NA/RNA binding <u>p</u>rotein of <u>43</u> kDa (TDP-43) mislocalizes from the nucleus to the cytoplasm (5–7). Misfolded and hyperphosphorylated TDP-43 forms inclusions in the cytoplasm of neurons in the brain and spinal cord of ALS patients (8). This TDP-43 proteinopathy is also found in the majority of FTD patients and over 50% of AD patients and 60% of PD patients (9, 10), indicating a broader role (mechanism?) of TDP-43 in the pathogenesis of multiple neurodegenerative diseases. A recent report suggests that accumulation of TDP-43 in limbic structures defines a new type of dementia (11). TDP-43 has a high propensity to aggregate due to its carboxyterminal prion-like domain (12, 13). Accordingly, prediction of prion-like and aggregation-prone proteins in humans by an algorithm ranks TDP-43’s propensity to aggregate as 69^th^ out of the entire human proteome (14).

Cells combat the accumulation of damaged or misfolded proteins by the action of molecular chaperones and heat shock proteins, which facilitate protein refolding or degradation to prevent the toxic consequences associated with protein misfolding (15). Hsp90 is a cytosolic molecular chaperone that is expressed at high levels under normal conditions and strongly induced by stress. It is highly conserved within eukaryotes and regulates many cellular pathways, such as cellular signaling, cell cycle control, and cell survival (16).

Hsp90 co-localizes with many disease-related protein aggregates, such as tau tangles and amyloid beta (Aβ) plaques in AD (17, 18). It has also been shown to inhibit tau and Aβ aggregation (19, 20), which may represent a defense mechanism. Yet, it has also been proposed that Hsp90 binds and stabilizes toxic aberrant protein species, leading to the toxic accumulation of these proteins and thereby contributing to neurodegeneration (18). For instance, epichaperomes, i.e. metastable chaperone networks stabilized by Hsp90, seem to promote the accumulation and toxicity of misfolded tau (21). Experiments using purified proteins have shown that Hsp90 has the capacity to both inhibit TDP-43 aggregation and to convert TDP-43 aggregates into more soluble species (22).

The binding and hydrolysis of ATP is essential for Hsp90 binding to client proteins. Co-chaperones, such as <u>a</u>ctivator of <u>H</u>sp90 <u>A</u>TPase protein <u>1</u> (Aha1), <u>c</u>ell <u>d</u>ivision <u>c</u>ycle protein <u>37</u> (Cdc37), and <u>st</u>ress inducible phosphoprotein <u>1</u> (Sti1, also known as STIP1 and <u>H</u>sp70/Hsp90 <u>o</u>rganizing <u>p</u>rotein, HOP in humans) inhibit (Sti1 and Cdc37) or stimulate (Aha1) Hsp90 ATPase activity and facilitate the folding or activation to specific client proteins (23).

Aha1 stimulates the ATPase activity of Hsp90 by up to twelve fold over its basal level (24, 25) and regulates the activation of Hsp90 client proteins (26). Cdc37 regulates the activity of protein kinases; it can act as a molecular chaperone independently of Hsp90 and its over expression can compensate for decreased Hsp90 function (27). Cdc37 together with Hsp90 participate in the nuclear localization of TDP-43, which is typically associated with reduced TDP-43 toxicity; the disruption of the Cdc37/Hps90 complex triggers degradation of TDP-43 via autophagy (28).

Unlike other Hsp90 co-chaperones, Sti1 can bind to both Hsp70 and Hsp90 simultaneously to allow for client protein transfer between these two major chaperone systems (29). Sti1 can function independently of Hsp90 and is involved in protein folding of protein kinases and the curing of yeast prions through interaction with Cdc37 (30, 31). Sti1 also interacts with many different aggregation prone proteins (32–34), including the mammalian prion protein (35).

Collectively, these studies document the capacity of Hsp90 and its co-chaperones to modulate protein misfolding, aggregation, and the ensuing toxicity. Although Hsp90 has been shown to interact with TDP-43, it remains unclear whether Hsp90 and its co-chaperones modulate TDP-43 toxicity. Here we report for the first time that Hsp90 and its co-chaperones, particularly Sti1, regulate TDP-43 aggregation and toxicity in yeast and mammalian cell models.

## Results

### Hsp90 regulates TDP-43 toxicity

We used well-established yeast models to explore how Hsp90 levels and its ATPase activity modulate TDP-43 proteinopathy and toxicity (13, 36–38). In a slight variation to previously published TDP-43 yeast models, we used TDP-43-GFP expressed from a low-copy plasmid (CEN) under control of the inducible galactose (GAL-1) promotor, to provide low level overexpression, which produces moderate growth defects (Fig 1A). This model allows us to study factors that can either enhance or suppress TDP-43 toxicity. We first sought to determine the effect of inhibiting Hsp90’s ATPase activity by treating yeast cells expressing TDP-43-GFP with two established Hsp90 inhibitors, radicicol and 17AAG (17-(Allylamino)-17-demethoxygeldanamycin) (39, 40). Growth assays on agar plates (Fig 1B) and liquid cultures (Fig 1C) document an increase in TDP-43 toxicity even at low concentrations of both Hsp90 inhibitors. Of note, these low concentrations of radicicol and 17AAG do not cause growth defects in wild type control cells or elicit the heat shock response (HSR) (41). Fluorescence microscopy (Fig 1D) shows no major difference in TDP-43 localization between untreated and treated cells with somewhat reduced cytoplasmic inclusions and more diffuse localization of TDP-43-GFP in radicicol- and 17AAG-treated cells compared to untreated cells.

**Figure 1:**
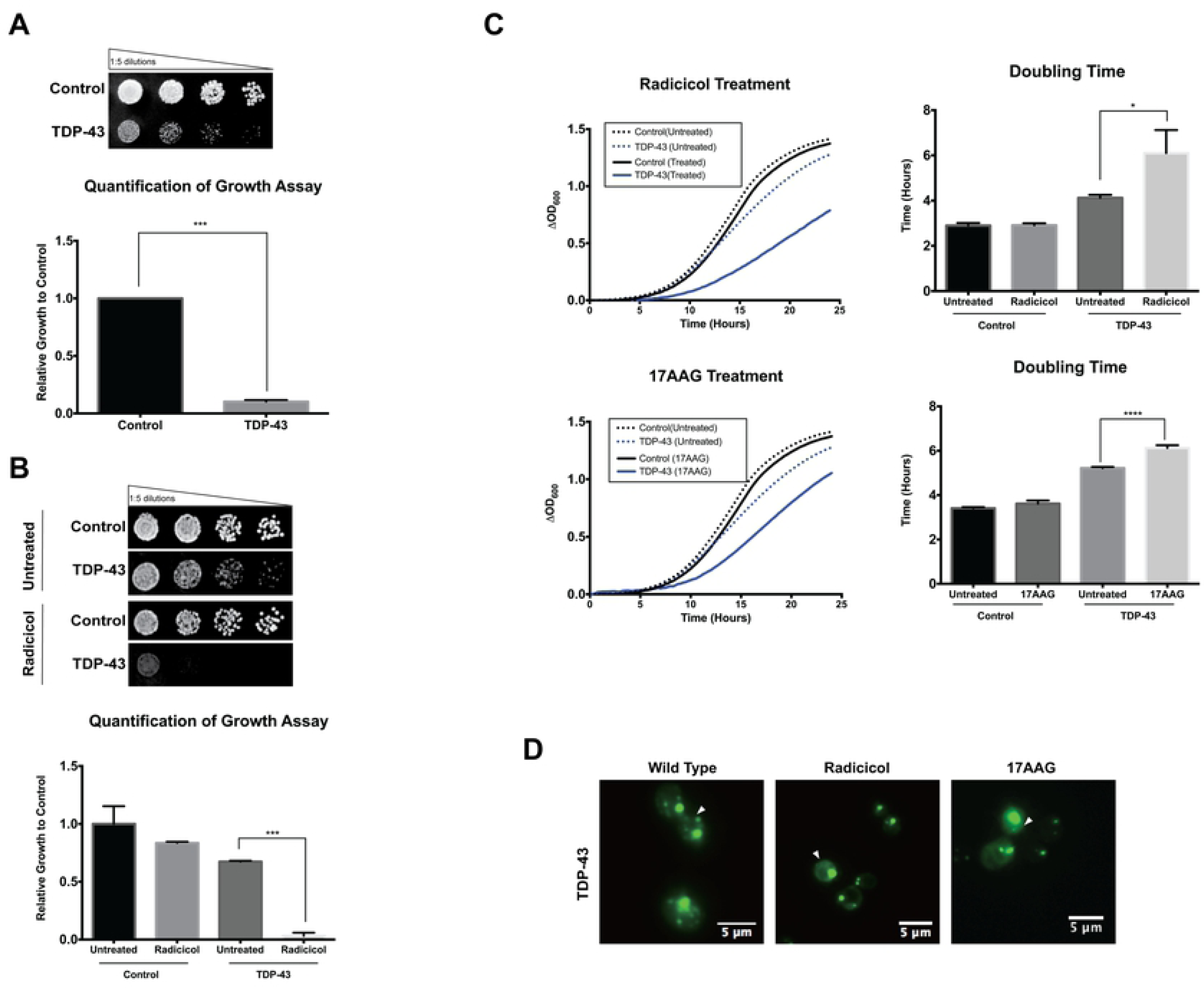
Inhibition of Hsp90’s ATPase activity increases TDP-43 toxicity and alters its localization. **A** Growth assay with wild type yeast cells expressing TDP-43-GFP or a vector control. Cells were spotted in four five-fold serial dilutions (from left to right) on plates that induce TDP-43-GFP. TDP-43 toxicity is inferred by the lack of growth in the TDP-43 lanes when compared to the vector control. **B** Growth assay of wild type yeast cells expressing TDP-43-GFP on control plates and plates containing 10 μg/mL radicicol (∼27.4 μM). **C** Growth curves of wild type yeast cells expressing TDP-43-GFP and cells bearing a vector control grown in the presence or absence of 10 μg/mL radicicol and 10 mM 17-AAG. The doubling time of the cells was calculated during the log phase of the growth curve. **D** Fluorescence microscopy of wild type yeast cells grown for 16 h in inducing medium in the presence or absence of 10 μg/mL radicicol and 10mM 17-AAG show nuclear localization of TDP-43 in Hsp90 inhibitor-treated cells.

If Hsp90 regulates TDP-43 toxicity, reducing cellular Hsp90 levels should reproduce the effects of pharmacological inhibition. Deletion of both Hsp90 alleles together in yeast (*Hsc82*, the constitutive allele and *Hsp82*, the stress induced allele) is lethal, whereas the deletion of either one allele alone does not result in adverse growth phenotypes under normal conditions (42). TDP-43-GFP was expressed in yeast cells that lack either one of the yeast *Hsp90* encoding genes (*hsc82Δ* or *hsp82Δ* yeast strains). Our results show an increase in TDP-43 toxicity in both Hsp90 deletion strains when compared to wild type controls (Fig 2A). Fluorescence microscopy did not reveal any striking differences in TDP-43-GFP localization and aggregation between wild type and either Hsp90 deletion strain (Fig 2B). Occasionally, Hsp90-depleted cells displayed a slight increase in smaller cytosolic puncta, which was more pronounced in the *hsc82Δ* than in the *hsp82Δ* strain. These results indicate that inhibiting Hsp90 ATPase activity or reducing cellular Hsp90 levels increases TDP-43 toxicity.

**Figure 2:**
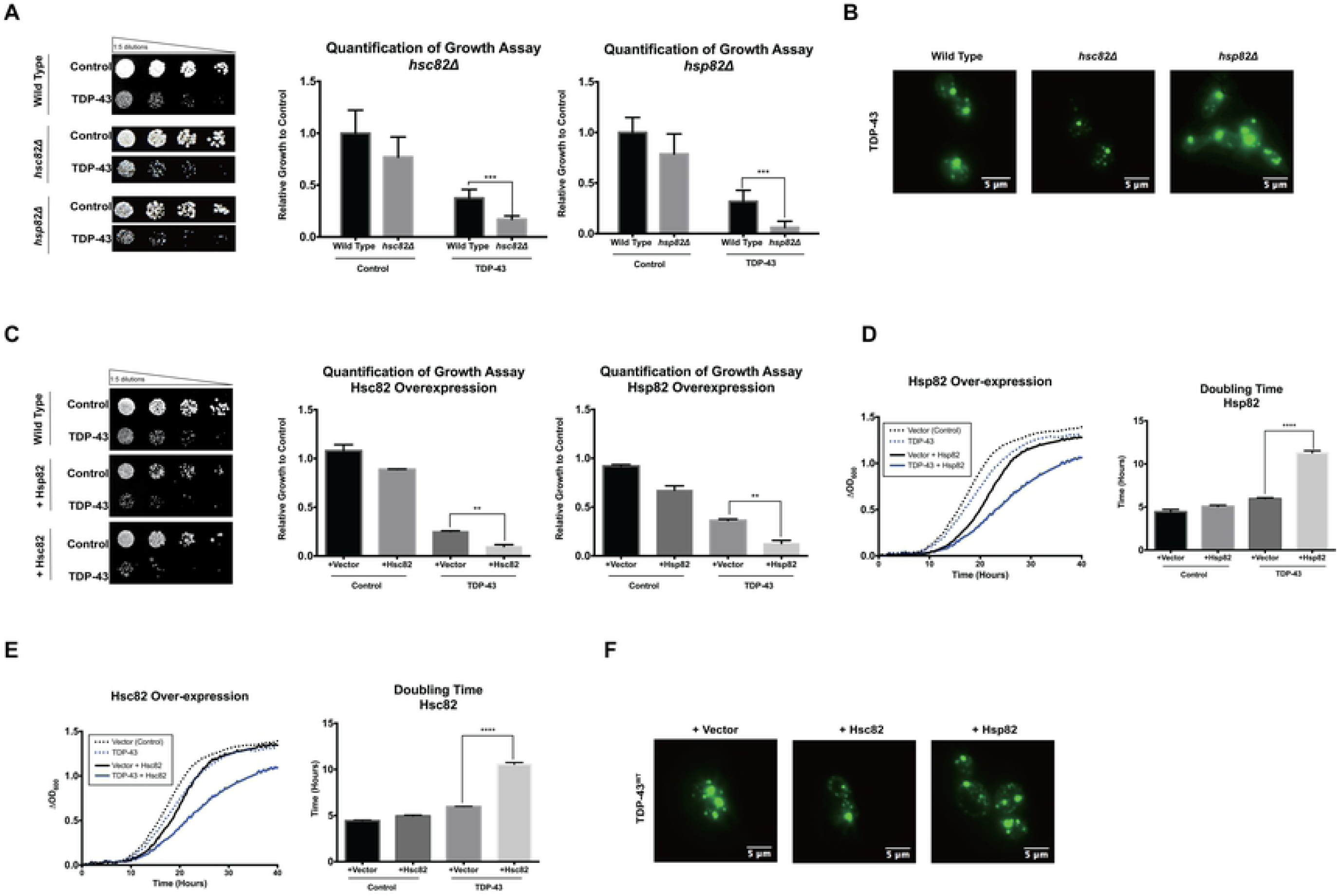
Decreasing and increasing Hsp90 levels exacerbate TDP-43 toxicity and alter its localization. **A** Growth assay and **B** fluorescence microscopy of wild type and Hsc82 or Hsp82 deleted cells expressing TDP-43-GFP. **C** Growth assay of wild type yeast cells over-expressing Hsc82 or Hsp82 that co-express TDP-43-GFP or a vector control. Liquid growth curves and doubling time of wild type yeast cells over-expressing. **D** Hsp82 and **E** Hsc82 that co-express TDP-43-GFP or a vector control. **F** Fluorescence microscopy of wild type yeast cells over-expressing Hsc82 or Hsp82 that co-express TDP-43-GFP.

### Over-expression of Hsp90 exacerbates TDP-43 toxicity

We next tested whether Hsp90 over-expression alters TDP-43 toxicity by over-expressing *Hsc82* or *Hsp82* using a high copy plasmid (2µ). When co-expressed with TDP-43-GFP, over-expression of *Hsc82* or *Hsp82* caused a substantial increase in TDP-43 toxicity (Fig 2C). This synthetic toxicity is especially apparent in liquid culture growth (Fig 2D-E), where the doubling times of *Hsp82* or *Hsc82* over-expressing cells that co-express TDP-43-GFP are significantly increased. Fluorescence microscopy in cells over-expressing either *Hsc82* or *Hsp82* are very similar to control cells with some cells showing slightly increased diffuse cytoplasmic localization of TDP-43-GFP (Fig 2F).

### Sti1, Aha1, and Cdc37 and TDP-43 toxicity

Given the robust effects of changing Hsp90 on increasing TDP-43 toxicity, we tested whether modulating Hsp90 co-chaperones could also affect TDP-43 aggregation and toxicity. We focused on three co-chaperones that have been previously implicated in regulating protein misfolding (16, 28, 43, 44). First, we tested the effect of deletion of *Sti1* or *Aha1* (*sti1Δ* or *aha1Δ* yeast strains respectively), and reduced expression of the essential *Cdc37* (Cdc37-DAmP yeast strain). We also included two essential genes, *Ess1* and *Sgt1*, in our studies to test whether reduced expression of these genes had an effect on TDP-43 toxicity. *Ess1* is an essential gene that encodes a peptidylprolyl cis/trans isomerase that plays a role in transcription regulation. *Sgt1* is an essential Hsp90 co-chaperone that has been implicated in kinetochore complex assembly and in neurodegeneration (45, 46).

Our results show that deletion of *Sti1*, which did not have any adverse effect on the growth of control cells, drastically exacerbated TDP-43 toxicity. In contrast, deletion of *Aha1* had no effect on TDP-43 toxicity (Fig 3A). Due to the innate toxicity associated with reduced expression of Cdc37, we chose a construct that expresses TDP-43 at a very low level and exhibits little to no toxicity when expressed in wild type cells to test for TDP-43 toxicity in the Cdc37-DAmP strain. We observed an increase in TDP-43 toxicity when TDP-43 is expressed in the Cdc37-DAmP strain (Fig 3B). However, it should be noted that reducing Cdc37 also affects the wild type strain, hence the effect of Cdc37 may not be selective to TDP-43. Synthetic toxicity is not seen in reduced expression of the other two essential Hsp90 co-chaperones, *Ess1* and *Sgt1* (S1A-B Fig).

**Figure 3:**
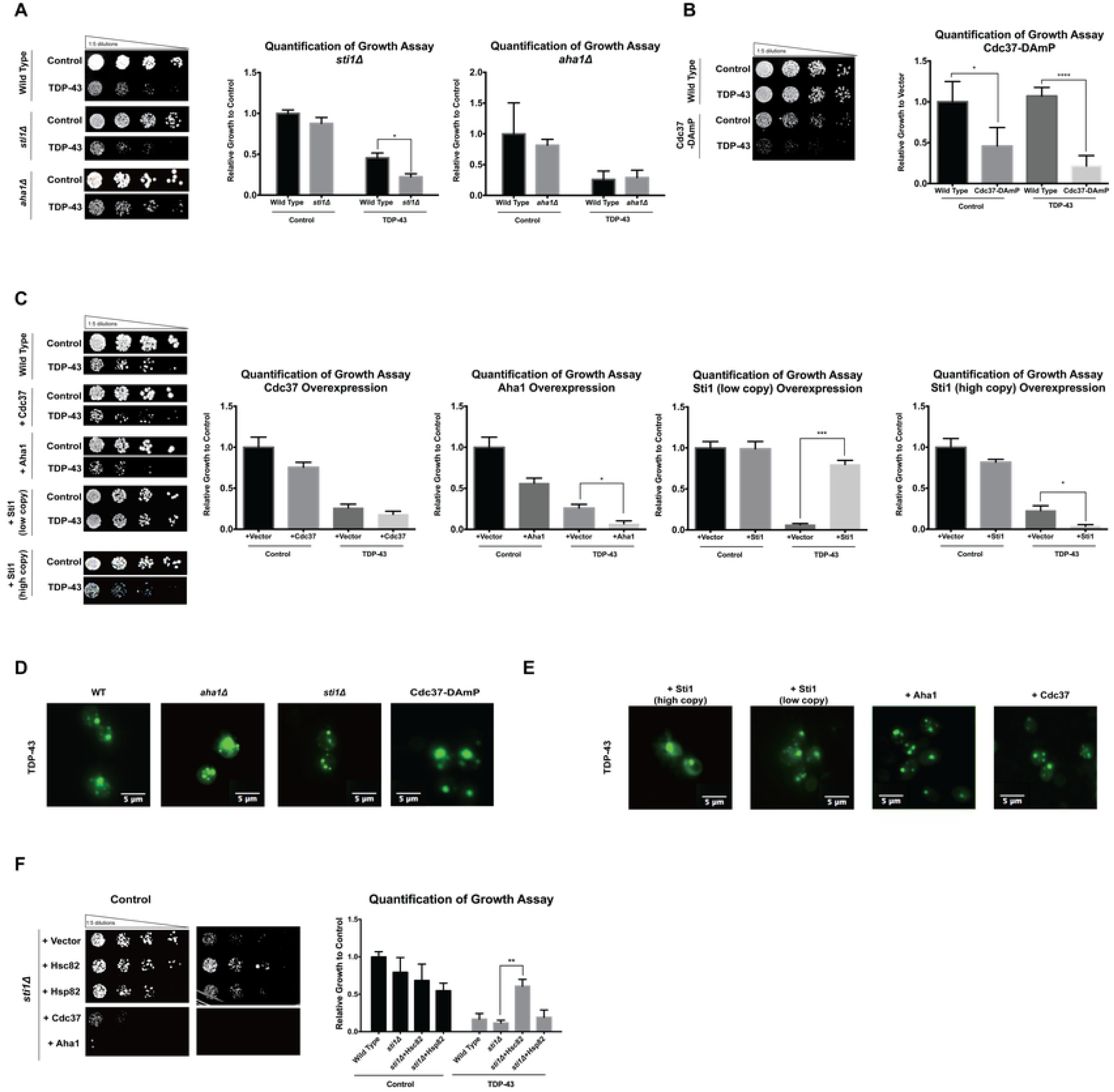
Hsp90 co-chaperones and TDP-43. Growth assay of wild type yeast cells and yeast cells deleted for **A** Sti1 or deleted for Aha1, and **B** reduced expression of Cdc37 expressing TDP-43-GFP or bearing a vector control. **C** Growth assay of wild type yeast cells moderately over-expressing (CEN) Sti1, Cdc37, Aha1 or highly over-expressing Sti1 (2μ) Sti1 that co-express TDP-43-GFP. Fluorescence microscopy of **D** wild type yeast cells and yeast cells deleted for Sti1, and reduced expression of Cdc37 and **E** wild type yeast cells moderately over-expressing (CEN) Sti1, Cdc37, Aha1 or highly over-expressing Sti1 (2μ) Sti1 that co-express TDP-43-GFP. **F** Growth assay of yeast cells deleted for Sti1 moderately over-expressing Hsc82, Hsp82, Cdc37, Aha1, that co-express TDP-43-GFP.

We then assessed how *Sti1, Aha1*, and *Cdc37* over-expression affects TDP-43 toxicity. Given the surprising effects of high Hsp90 over-expression, we used two different expression systems—high over-expression from high copy plasmids (2µ) and moderate over-expression from low copy plasmids (CEN). High over-expression of *Sti1* resulted in a slight reduction of growth in control cells, but high over-expression of *Aha1* and especially *Cdc37* showed significant reduced growth in control cells (Fig 3C, S2A Fig). In TDP-43-expressing cells, high over-expression of these co-chaperones produced an increase in toxic phenotypes (Fig 3C, S2A Fig). However, moderate over-expression of *Sti1* showed no toxicity, whereas moderate over expression of *Aha1* or *Cdc37* was toxic (Fig 3C). This increased toxicity is also seen when TDP-43 is co-expressed with *Aha1* and *Cdc37* (Fig 3C). By contrast, moderate *Sti1* over-expression rescued TDP-43 toxicity (Fig 3C). Fluorescence microscopy of TDP-43-GFP in the *sti1Δ* strain showed an increase in cytosolic aggregates that appear to be smaller in size, while TDP-43-GFP localization in the *aha1Δ* strain and the strain expressing reduced levels of *Cdc37* were indiscernible from wild type (Fig 3D). Notably, moderate over-expression of *Sti1* increase diffuse cytosolic TDP-43, whereas high over-expression shows more inclusions of TDP-43 (Fig 3E). Moderate *Aha1* and *Cdc37* over-expression did not produce obvious changes in TDP-43-GFP localization when compared to control cells (Fig 3E). Conversely, high over-expression of *Aha1* and *Cdc37* both increased diffuse cytosolic TDP-43-GFP localization (S2B Fig). Our results show that the Hsp90 co-chaperone *Sti1* and to a lesser extent *Aha1* or *Cdc37*, alter TDP-43 toxicity and its localization and aggregation. Of note, the effects of *Sti1* are dosage dependent concluded from the results showing that moderate *Sti1* over-expression reduced TDP-43 toxicity, whereas high *Sti1* over-expression exacerbated TDP-43 toxicity.

Hsp90 and *Sti1* deletion show synthetic lethality (47) and there is ample genetic evidence for their co-regulation. We therefore sought to investigate how the over-expression of Hsp90, *Aha1*, or *Cdc37* in *sti1Δ* yeast strains controls TDP-43 toxicity (Fig 3F), to determine whether overexpression of other molecular chaperones or co-chaperones can compensate for the loss of *Sti1*. To this end, we highly over-expressed Hsp90 (both *Hsc82* and *Hsp82*) in the *sti1Δ* strain that also expressed TDP-43-GFP. The growth assay shown in Fig 4 indicates that high over-expression of *Hsc82* suppressed TDP-43 toxicity in the *sti1Δ* strain, whereas high over-expression of both *Aha1* and *Cdc37* strongly enhanced TDP-43 toxicity in the *sti1Δ* strain (S2C Fig). These results reveal that TDP-43 toxicity is regulated by multiple genetic interactions of Hsp90 and that an imbalance of Hsp90 and its co-chaperones exacerbates TDP-43 toxicity.

**Figure 4:**
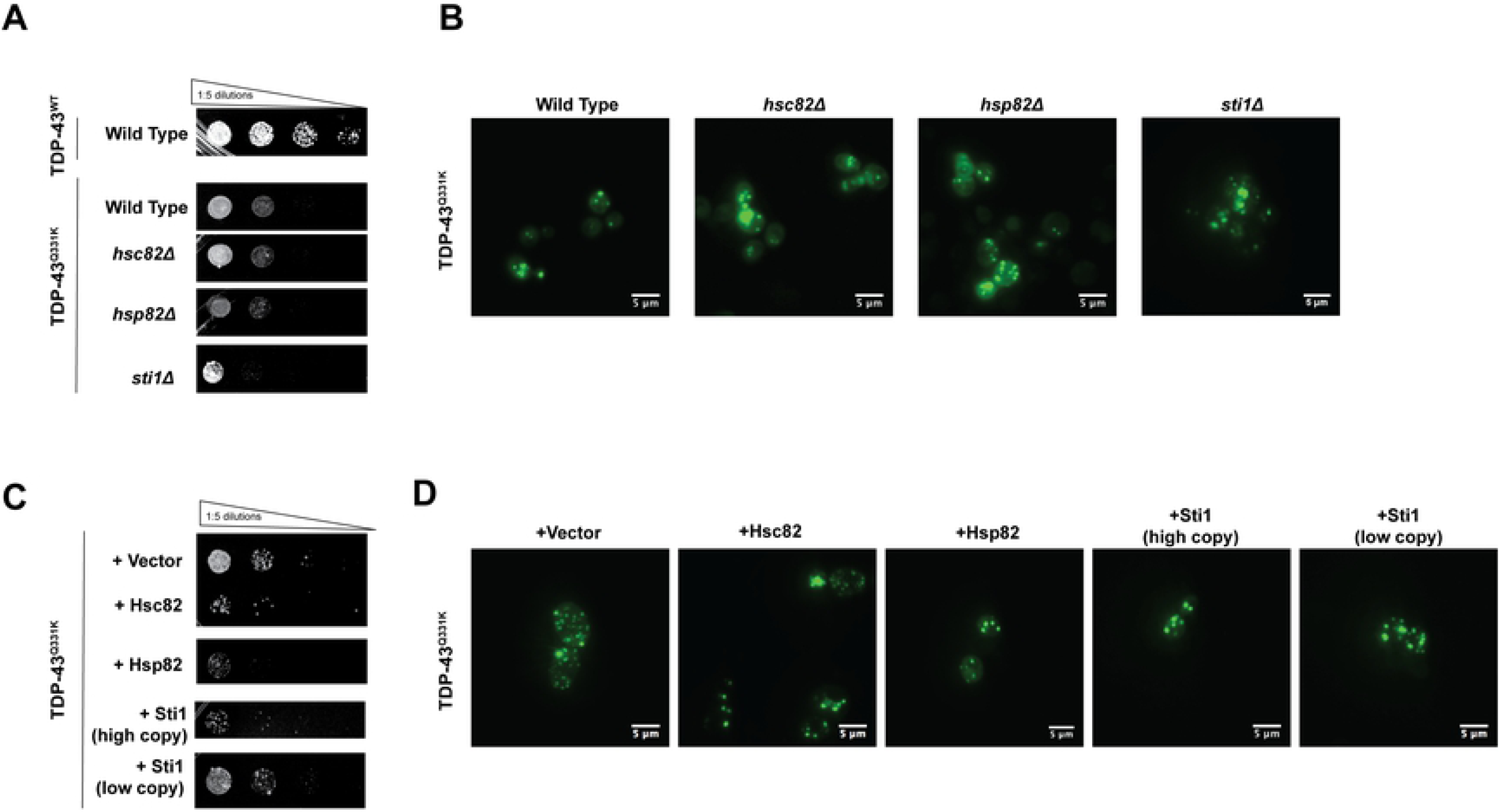
Hsp90 and Sti1 do not alter TDP-43^Q331K^ toxicity and localization. **A** Growth assay and **B** fluorescence microscopy of wild type and cells deleted for Hsc82, Hsp82, or Sti1 expressing TDP-43^Q331K^-GFP or bearing a vector control. **C** Growth assay and **D** fluorescence microscopy of wild type cells highly (2μ) over-expressing Hsc82 and Hsp82 or moderately (CEN) or highly (2μ) over-expressing Sti co-expressing TDP-43^Q331K^-GFP or bearing a vector control.

We also tested a suite of Sti1 variants bearing point mutations in and truncations of key domains (S3 Fig), to determine whether different functional domains of Sti1, such as its TPR domains, are required for its interaction with TDP-43. None of these variants exacerbated or reduced TDP-43 toxicity indicating that only full-length wild type Sti1 protein can modulate TDP-43 in this manner.

### Hsp90 and its co-chaperones have no effect on TDP-43 Q331K

Mutations in the gene encoding TDP-43 are associated with familial forms of ALS (fALS). The majority of fALS TDP-43 mutations result in amino acid substitutions in the prion-like domain at the carboxy-terminus of TDP-43 (48). These mutations are thought to result in increased TDP-43 proteinopathy and toxicity (13). Yeast models, e.g. for the fALS mutant TDP-43 Q331K, indeed show increased aggregation and toxicity (13). Our experiments document that unlike wild-type TDP-43, manipulating Hsp90 and *Sti1* activity in yeast had no significant effect on TDP-43 Q331K toxicity and aggregate localization and solubility (Fig 4A-D), suggesting that the conformation assumed by this mutant makes it independent of the Hsp90 chaperone machinery in yeast.

### Higher Hsp90 levels increase TDP-43 protein levels

Our next experiments aimed to decipher the cellular mechanism by which Hsp90 and its co-chaperone affects TDP-43. We asked whether changes in TDP-43 toxicity and localization in yeast cells expressing different levels of Hsp90 and *Sti1, Aha1*, or *Cdc37* may be the result of altered TDP-43 protein levels. We thus monitored TDP-43-GFP levels in yeast cells bearing deletions or over-expression of Hsp90 and its co-chaperones. Our Western blot analyses reveal that Hsp90 over-expression (*Hsp82* or *Hsc82*) increased TDP-43 levels at steady-state, whereas TDP-43 remained unaltered in the *hsc82Δ* and *hsc82Δ* yeast strains (Fig 5A and 5B). Interestingly, TDP-43 protein levels were reduced in the *sti1Δ* strain even though TDP-43 toxicity was increased (Fig 5C). Deletion of *Aha1*, the moderate and high over-expression of *Sti1, Aha1*, and *Cdc37* had no effect on TDP-43 protein levels (S4 Fig). These results indicate that increased Hsp90 levels increases TDP-43 protein levels and thus exacerbated its toxicity. By contrast, decreasing Hsp90 did not alter TDP-43 levels. In addition, while over-expression of *Sti1* decreases TDP-43 toxicity, these effects were not associated with altered TDP-43 protein levels.

**Figure 5:**
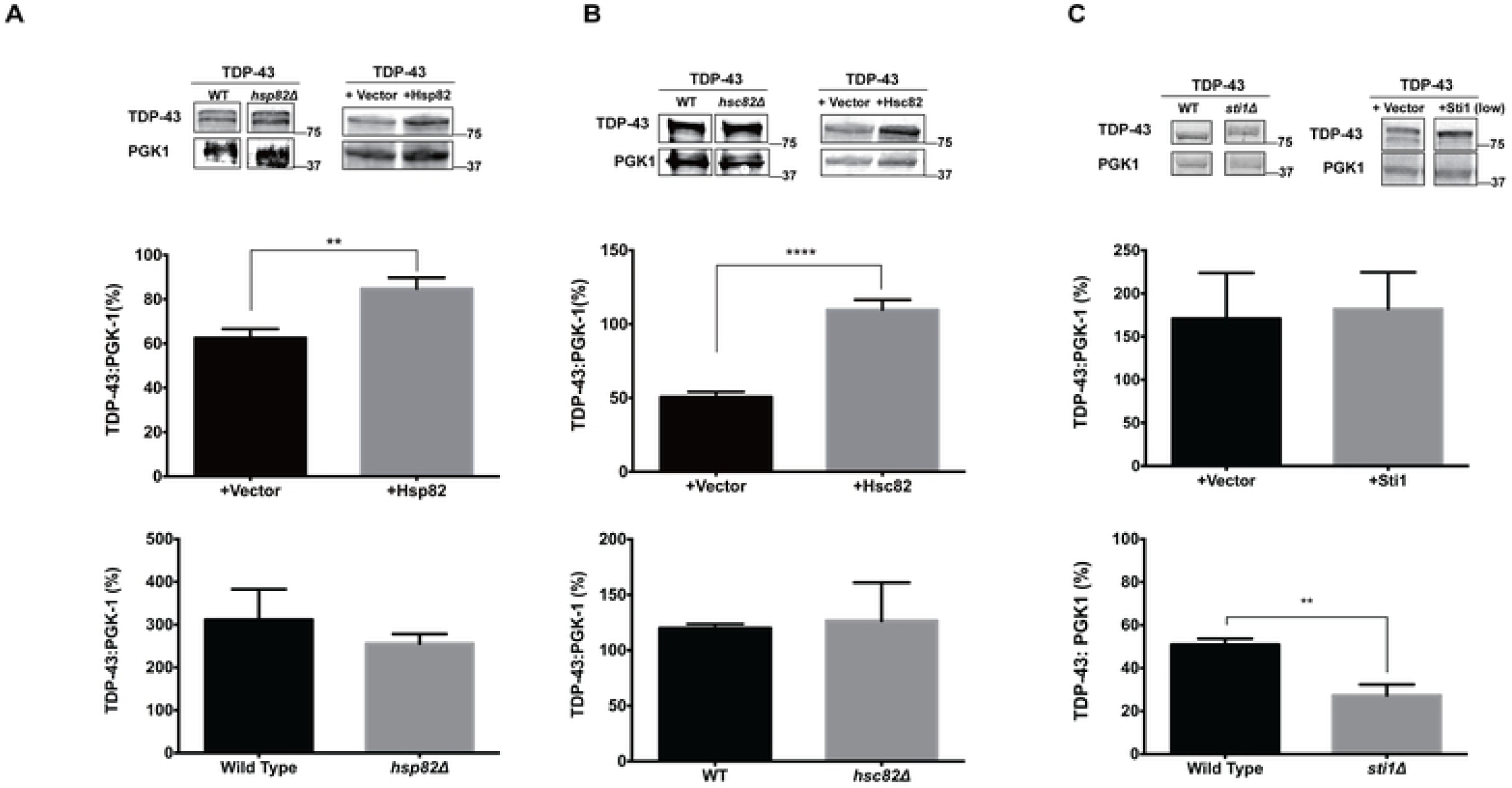
Over-expressed Hsp90 increases TDP-43 levels. Immunoblots and their quantitative analysis prepared with protein lysates from yeast cells over-expressing or deleted **A** Hsp82, **B** Hsp82, C **Sti1** or bearing vector controls co-expressing TDP-43-GFP probed with an anti-GFP antibody. Pgk1 served as a loading control. The quantitative analysis represents the average of three independent experiments. Error bars represent standard errors of the mean.

### Sti1 modulates TDP-43 solubility

The aggregation and mislocalization of TDP-43 from the nucleus to the cytoplasm is a central pathological hallmark of ALS and other TDP-43-associated neurodegenerative diseases (8, 49) that can be faithfully recapitulated in yeast models. By contrast, the biophysical properties of these TDP-43 inclusions seems more variable. TDP-43 can form both detergent- and denaturing-resistant aggregates and more soluble amorphous protein deposits (50). These different biophysical properties may depend on the experimental system, expression levels, and growth conditions. Therefore, following a previously established protocol (51, 52), we devised a sedimentation assay to assess TDP-43 solubility in the presence of mild detergents. These sedimentation assays did not reveal any difference in solubility under conditions of increased Hsp90 expression levels (Fig 6A and 6B). Neither the over-expression of *Aha1* nor *Cdc37* showed any changes in TDP-43 solubility in sedimentation experiments (Fig 6C and 6D). Comparatively, *sti1Δ* strains showed an increase in the relative proportion of insoluble TDP-43 (Fig 6E). Both moderate and high over-expression of *Sti1* also resulted in increased levels of soluble TDP-43 (Fig 6F and 6G).

**Figure 6:**
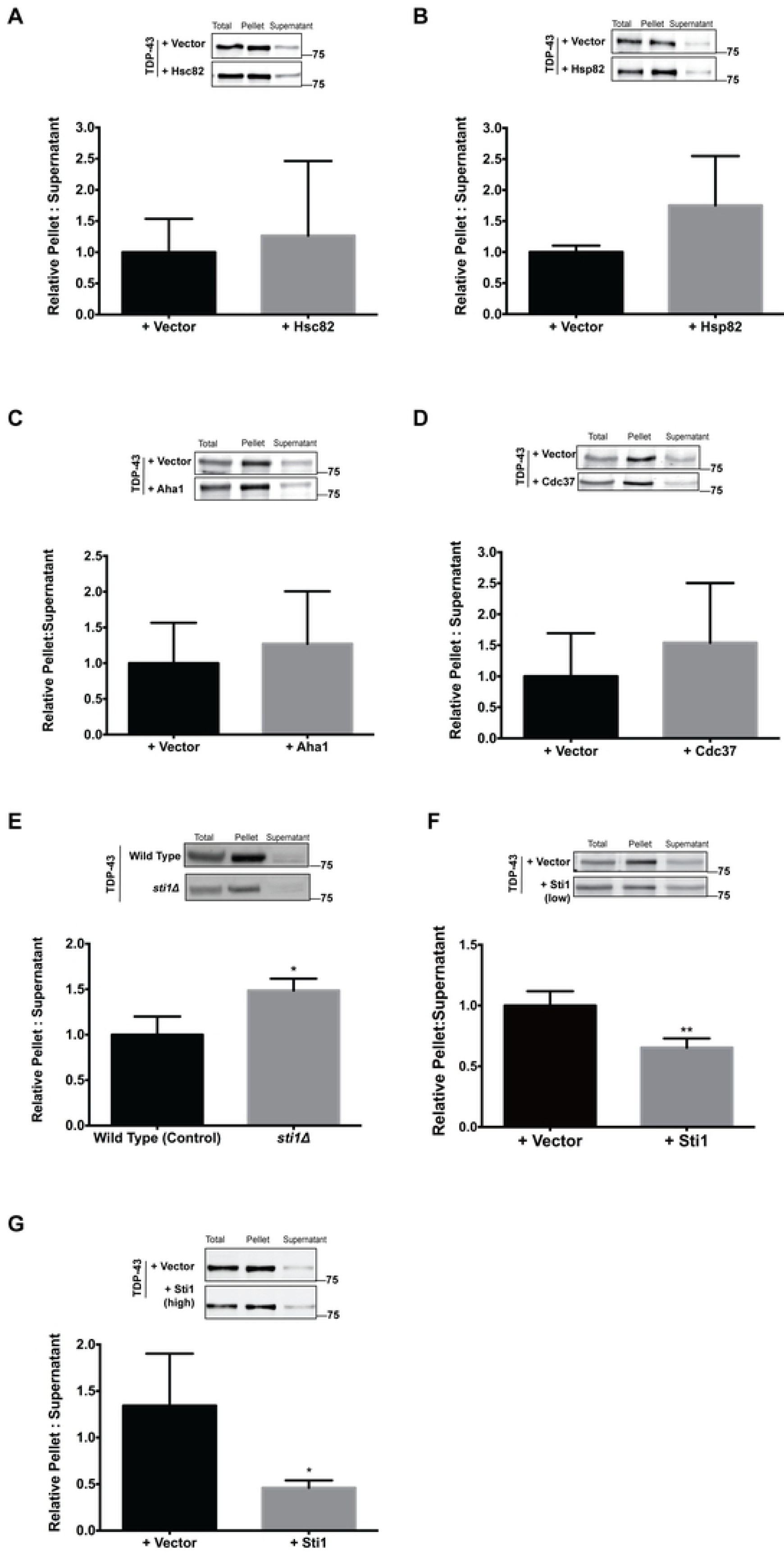
Sti1 alters TDP-43 solubility. Sedimentation analysis of yeast cells over-expressing **A** Hsc82 (high expression, 2μ plasmid) **B** Hsp82 (high expression, 2μ plasmid), **C** Aha (high expression, 2μ plasmid), **D** Cdc37 (high expression, 2μ plasmid), **E** Sti1 deletion, **F** Sti1 (low expression, CEN plasmid), **G** Sti1 (low expression, CEN plasmid), show that TDP-43 solubility is altered when co-expressed with Sti1. Blots were probed with an anti-GFP antibody. Pgk1 served as a loading control. The quantitative analysis represents the average of three independent experiments. The vector control samples served as references for the relative changes in the samples with altered Sti1 levels. Error bars represent standard errors of the mean.

### Sti1 modulates TDP-43 proteinopathy and toxicity in mammalian systems

Next, we examined the interaction between Sti1 and TDP-43 in mammalian cell lines. For these studies we first employed HEK 293 and HeLa cells (S5 Fig and S6 Fig). Yet, our experiments revealed severe limitations in both cell lines. The overwhelming majority of cells showed exclusively nuclear TDP-43 localization with little or no toxicity (S5B Fig and S6B Fig). We speculate that the constitutive and high induction of the heat shock response in cancerous cells with high-division rates (53), such as HEK293 and HeLa cells, results in high steady-state levels of many molecular chaperones, rendering them unusually resilient towards protein misfolding toxicity. We therefore employ two neuronal cell lines for our experiments, mouse Septal Neuronal (SN56) cells and mouse neuroblastoma Neuro2N (N2a) cells, both of which express neuronal marker proteins and have served in previous protein misfolding studies, including central hallmarks of TDP-43 proteinopathy for our experiments.

Using CRISPR-Cas9 to knockout Sti1, we engineered SN56 cells that lacked Sti1 (Sti1-KO) (54). These Sti1-KO cells showed a markedly reduced level of endogenous TDP-43 protein compared to wild type SN56 cells, without any change in TDP-43 mRNA levels (Fig 7A). In addition, tissue from Sti1-KO mouse embryos (55) showed that endogenous TDP-43 levels were found to be reduced in Sti1-KO embryos (Fig 7B and 7C). These results suggest a conserved role for Sti1 in modulating TDP-43 levels in yeast, mammalian cells, and mouse embryos.

**Figure 7:**
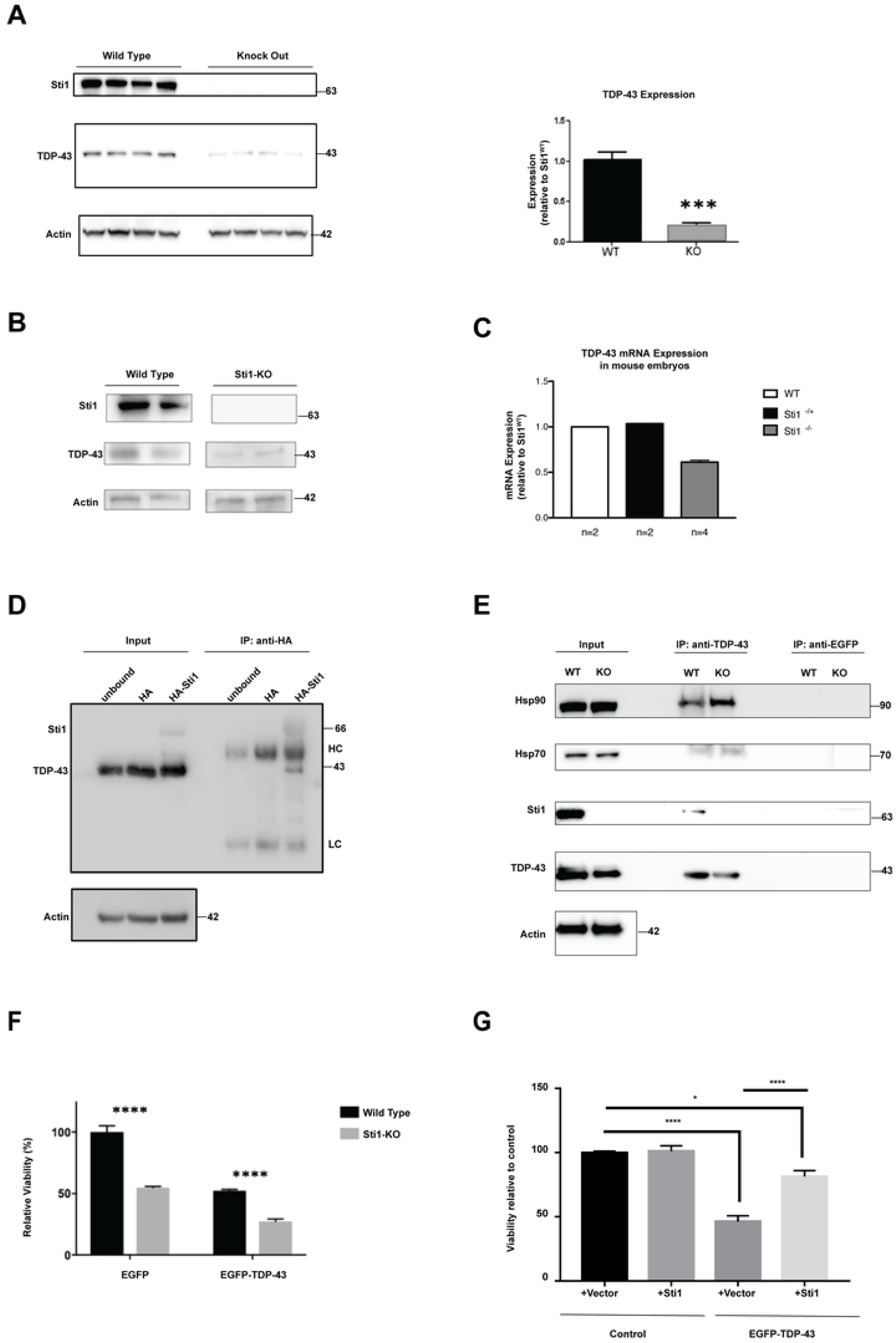
Sti1 interacts with TDP-43 in mammalian cells. **A** CRISPR-Cas9 was used to eliminate Sti1 expression in neuronal cell cultures (SN56). Proteins extracted from wild type and Sti1 KO cell cultures were resolved using SDS-PAGE and blotted with antibodies against Sti1 and TDP-43 and quantified. **B** Western blots were performed with Sti1 wildtype and KO embryos tissue lysates and TDP-43 was found to be reduced in Sti1-KO embryos. n=2 **C** TDP-43 mRNA expression was reduced in Sti1^-/-^ KO mouse embryos but unchanged in Sti1^+/-^ KO mouse embryos. **D** HEK293 cells were transfected with HA-Sti1, HA-empty vector, or non-transfected and anti-HA agarose beads were used to precipitate HA-STIP1. Co-IP proteins were resolved by SDS-PAGE, transferred to a membrane and blotted with an antibody against TDP-43 or HA. HC and LC are heavy and light chain of antibodies respectively. **E** SN56 cells were used to co-immunoprecipitate Hsp90, Hsp70 and Sti1 with endogenous TDP-43. Input represents total cell extracts used for immunoprecipitation. IP TDP-43 lanes represent endogenous TDP-43 immunoprecipitated extracts. Control (IP GFP) lanes represent extracts that were immunoprecipitated with antibody anti-GFP. For each of the three conditions both STIP1-WT (WT) and STIP1-KO SN56 cells (KO) were used. **F** SN56 WT and Sti1 KO cells were transfected with GFP-Empty vector, GFP-WT-TDP-43. Luciferase assay was performed to measure cellular viability. **G** SN56 WT cells were transiently transfected with EGFP vector or EGFP-TDP-43 and co-expressed with either a HA-empty vector control or HA-Sti1. The cells were differentiated and then monitored for TDP-43 toxicity through a luciferase assay.

### Hsp70, Hsp90, and Sti1 form a complex with TDP-43 and regulate its toxicity in neuronal cell

Hsp90 and Hsp70 can form protein complexes with TDP-43 (28, 56, 57). To test the possibility that Sti1 can also form a complex with TDP-43, we immunoprecipitated HA-Sti1 and tested for specific interaction with TDP-43 in HEK cells (Fig 7D). The results show that Sti1 precipitates TDP-43. We then performed the reciprocal experiment and immunoprecipitated GFP-TDP-43 from wild type SN56 cells and tested for co-immunoprecipitation of a complex containing Sti1, Hsp70, and Hsp90 (Fig 7E). Indeed, we observed that TDP-43 is part of a complex with Sti1, Hsp70 and Hsp90, suggesting that the regulation of TDP-43 misfolding may depend on the formation of a complex with Hsp90, Hsp70 and Sti1. We repeated these experiments in Sti1-KO SN56 cells and tested whether endogenous Sti1 is required for the interaction of TDP-43 with both Hsp90 and Hsp70 (Fig 7E). Interesting, even in the absence of Sti1, TDP-43 could still precipitate both Hsp90 and Hsp70, suggesting that these molecular chaperones can interact with TDP-43 in the absence of Sti1. In sum, our results obtained with SN56 cells recapitulated key results from yeast and demonstrated a direct biochemical interaction between Sti1 and TDP-43. Importantly, in STI1-KO cells, transfection of TDP-43 increased cell death and toxicity (Fig. 7F). Conversely, increased expression of Sti1 in SN56 cells decreased TDP-43 toxicity (Fig. 7G).

Using a second cell model, we performed experiments in partially differentiated N2a cells to confirm these results in a second cell line and further evaluate the stoichiometry of Sti1 expression levels. These cells, particularly following serum withdrawal and limited glucose supply (see Material and Methods for details), quantitatively recapitulated TDP-43 proteinopathy and toxicity (Fig 8). We transiently co-transfected N2a cells with TDP-43 and different amounts of mouse Sti1 plasmids to produce varying expression levels of Sti1 (1/2x, 1x, and 2x, Fig 8). We monitored TDP-43 toxicity and found that, in parallel to our result in yeast and SN56 cells, moderate levels of Sti1 over-expression resulted in reduced TDP-43 toxicity. Yet unlike our yeast results, high Sti1 over-expression did not alter TDP-43 toxicity. Immunofluorescence microscopy revealed that moderate Sti1 over-expression in N2a cells resulted in an increase in nuclear localization of TDP-43 (Fig 8B, TDP-43 + Sti1 right two cells), whereas high Sti1 over-expression produces an increased number of small cytoplasmic puncta (Figure 8B, TDP-43 + Sti1 left cell). Consistent with our results in yeast, our sedimentation assays revealed increased TDP-43 solubility in the presence of moderate over-expression of Sti1. Lastly, we also confirmed that both TDP-43 and Hsp90 are part of a complex with Sti1 by co-immunoprecipitation in the N2a cells (Fig 8D).

**Figure 8:**
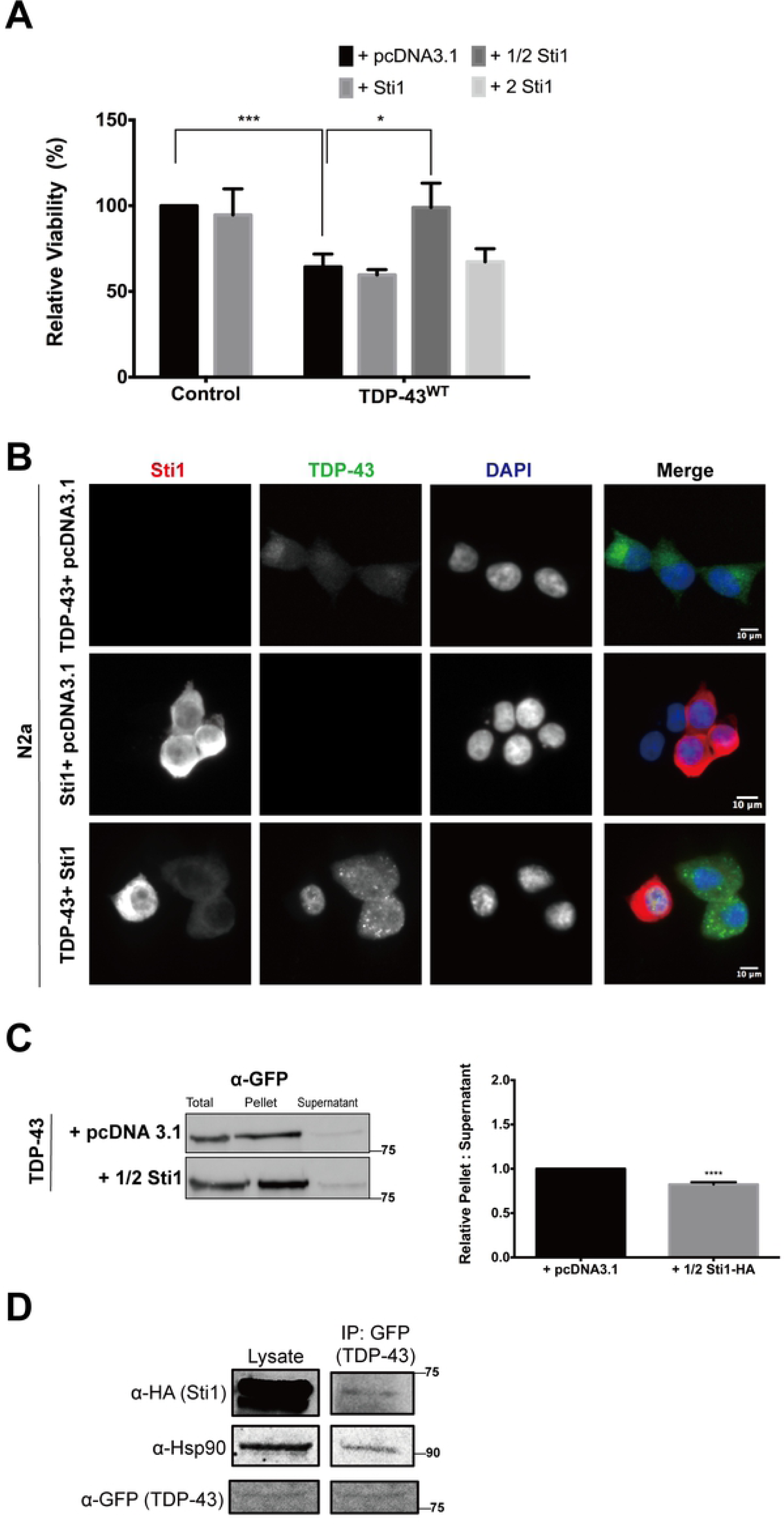
Sti1 modulates TDP-43 toxicity, localization, and solubility in N2a cells. **A** N2a cells were transiently transfected with a vector for expression of TDP-43-GFP together with a vector control or either equal amounts of Sti1 tagged with an HA epitope, or half (1/2x) and double (2x) the amount. The cells were partially differentiated and then monitored for TDP-43 toxicity through a luciferase assay. **B** Immunofluorescence microscopy of cells processed as described under (A) using GFP fluorescence and an anti-HA antibody for the detection of Sti1-HA. **C** Sedimentation assay of N2a cells process as described under (A) with the lower concentration (1/2x) of the Sti1-HA expression plasmid. The immunoblots (left panel) were probed with an anti-GFP antibody. The quantification (right panel) was derived from data of three independent experiments. Error bars represent standard errors of the mean. **D** Co-immunoprecipitation of Sti1 and Hsp90 with TDP-43 prepared from N2a protein lysates processed as described under (A).

## Discussion

Hsp90 and its co-chaperones have been suggested to have a highly select suite of client proteins, such as hormone receptors, kinases, and other signaling molecules (58, 59). Rather than regulating their de novo folding and preventing or reversing protein misfolding, Hsp90 and its co-chaperone supposedly regulate a final step in the maturation of its clients, which is essential for attaining their active conformation (44). Yet, Hsp90 and Sti1 also modulate the aggregation and toxicity of polyglutamine expansion proteins (60), tau (61) Aβ (34) and other misfolded proteins. In addition, we have shown increased expression levels of Sti1 in human AD brains and AD mouse models and that in vitro Sti1 decreases the toxicity of Aβ oligomers in cultured neurons and brain slices (62), whereas in vivo it favors amyloid aggregation (54). Our results suggest that the Hsp90 co-chaperone, Sti1, has critical roles in the regulation of TDP-43 misfolding and the ensuing toxicity, possibly by connecting the Hsp90 and Hsp70 chaperone systems to misfolded proteins (54). TDP-43 toxicity has important implications not only to ALS, but also to FTD, AD and possibly other neurodegenerative diseases. Our work in combination with published work thus contributes to an emerging view of a crucial role of Hsp90 and its co-chaperones in protein homoeostasis in the context of protein misfolding associated with neurodegenerative diseases (54, 63).

We find that decreased Hsp90 chaperone activity and decreased levels of Sti1 increase TDP-43 toxicity. This action of Sti1 seems to be selective. Altered levels of another Hsp90 co-chaperone, Aha1, does not seem to affect TDP-43 toxicity, whereas increased Cdc37 and Hsp90 expression exacerbates TDP-43 toxicity. Other essential Hsp90-associated genes in yeast also does not affect TDP-43 toxicity. Notably, moderate over-expression of Sti1 reduces TDP-43 toxicity, whereas high levels of Sti1 expression and its deletion increases TDP-43 toxicity. Moreover, we found that Sti1 forms a physical complex with TDP-43, Hsp90 and Hsp70 and that it regulates TDP-43 toxicity in a dose-dependent manner. Our results suggest that TDP-43 toxicity depends on properly balanced function of Hsp90 and its co-chaperones, particularly Sti1, and that disturbances of this balance sensitizes cells to TDP-43 toxicity. Changes in Sti1 levels can regulate the levels of classical clients of Hsp90 (54) and disturbances on Sti1 can contribute to abnormal chaperone function (35). Sti1 is also a critical node, which can disrupt epichaperomes, abnormal complexes of chaperone proteins that present increased connectivity due to protein misfolding (21). Remarkably, TDP-43 seems to behave as a client of Hsp90 to maintain its stability in cells.

Both, reduction of Hsp90 levels and, even more so, reduction of Sti1 levels, decreases TDP-43 protein. This result was observed in yeast, cultured cells and tissue of knockout Sti1 mice, suggesting that these interactions are conserved. It should be noted that TDP-43 has important developmental roles in regulating mRNA metabolism (12), and it is possible that the Hsp70 and Hsp90 chaperones are required for these physiological functions. Indeed, both Hsp90 (64) and Sti1 (65) have been found to have nuclear functions. For example, Hsp90 and Sti1 regulate the generation of Piwi microRNAs and transposable elements (66). Whether TDP-43 regulation of mRNA and microRNA can be regulated by molecular chaperones remains to be established.

All manipulations of the levels of Hsp90 and its co-chaperones that exacerbate TDP-43 toxicity in our experiments are associated with increased insolubility and altered cytosolic localization of TDP-43, either in multiple small inclusions, diffusely distributed protein, or both. By contrast, the protective moderate over-expression of Sti1 results in more soluble TDP-43 with increased nuclear localization, i.e. in partial reversal of TDP-43 proteinopathy and increased TDP-43 solubility. These results indicate that Sti1, possibly in concert with Hsp90 and Hsp70, may stabilize TDP-43 conformations that reduce TDP-43 proteinopathy and toxicity. Of note, while Sti1 can recruit multiple proteins to Hsp90 complexes, we show that TDP-43 can interact with both Hsp90 and Hsp70 independent of Sti1, indicating that these chaperones may regulate TDP-43 misfolding both together with Sti1 and without it. Our experiments in yeast show that in cells lacking Sti1, overexpression of Hsp82 prevents TDP-43 toxicity, a contrasting result when compared to wild-types cells in which overexpression of Hsp82 is toxic.

Given that Hsp90, Hsp70, and Sti1 form a complex with TDP-43 we cannot dismiss the possibility that misfolded and aggregated TDP-43 sequesters Hsp90 and its co-chaperones preventing them from properly performing their many essential functions, which can indirectly contribute to TDP-43 toxicity. This is less likely for the highly expressed Hsp90, but quite plausible for all co-chaperones, which are expressed at much lower levels (67). Finally, our data support the notion that a strong interaction between Hsp90 and some of its co-chaperones increase TDP-43 toxicity by accelerating TDP-43 export from the nucleus or stabilizing toxic TDP-43 conformations in the cytoplasm. All these different possible mechanisms are by no means mutually exclusive and may act together in regulating TDP-43 misfolding and toxicity.

Our data clearly does not provide the basis for a simplistic model whereby increased Hsp90 and its co-chaperone activity ameliorate TDP-43 toxicity and their reduced function exacerbate it. Rather, we show that the interaction between Hsp90, its co-chaperones, and TDP-43 is much more complex and depends on specific co-chaperones and their expression levels. This complexity plausibly originates from the multitude of different cellular functions of Hsp90 and the specific biophysical properties, cellular localization, and function of the misfolded protein, and the affected neuronal cell types. Future in vitro studies using purified protein and studies in animal models can help to clarify the exact mechanism by which Hsp90 and its co-chaperones’ regulation of TDP-43 proteinopathy and toxicity.

## Materials and Methods

### Materials

#### Chemicals

Radicicol, 17-AAG, and AZC were purchased from Sigma

#### Yeast strains

Yeast strains BY 4741 (MAT α his3Δ1 leu2Δ0 lys2Δ0 ura3Δ0) and W303 (MAT a leu2-3,112 trp1-1 can1-100 ura3-1 ade2-1 his3-11,15} [phi^+^]) were used in this study. Yeast deletion strains were obtained from the Saccharomyces Genome Deletion Project (68).

#### Yeast Media

Yeast-peptone-dextrose (YPD) rich media (10 g/L yeast extract, 20 g/L peptone, and 20 g/L dextrose) and selective dextrose (SD) media (2% glucose, 1X yeast nitrogen base (YNB), 6 g/L l-isoleucine, 2 g/L L-arginine, 4g/L L-lysine HCl, 6 g/L L-phenylalanine, 1 g/L L-threonine, and 1g/L L-methionine) in either liquid media or agar plates (20g/L) were used to grow yeast cells. SD media was supplemented with 4 different amino acids (4g/L L-tryptophan, 6g/L L-leucine, 2 g/L L-histidine-monohydrate) depending on the selectivity maker of the plasmid. 2% galactose or 2% galactose plus 2% raffinose was used instead of glucose as a carbon source to make selective galactose (SGal) and selective galactose raffinose (SGal Raf) media respectively, for induction of gene expression from plasmid with the GAL1 promoter.

#### Mammalian cell lines and media

SN65, N2a, HEK293T, and HeLa cell lines were used. Cells were grown in Dulbecco’s Modified Eagle Medium (DMEM, Corning) with 4.5 g/L and supplemented with 10% fetal bovine serum (FBS, Gibco), 1X penicillin-streptomycin solution (Corning), and 1X glutamine (Sigma Aldrich) at 37°C with ∼5% CO2. N2a cells were induced to differentiate in low glucose (1% glucose) and low serum (1% FBS) medium incubated for a minimum of 24 h (69). SN56 cells were differentiated as previously described (70). Differentiation was induced by incubating SN56 cells in complete median containing 1mM of dbcAMP (Sigma-Aldrich), a cAMP analog, for 48 hours.

#### Mouse Embryo

Wild type and Sti1^-/-^ mouse embryos were developed and collected as previously described (55). Briefly, heterozygous (Sti1^+/-^) females were super-ovulated and mated with heterozygous males. Super-ovulation was induced by injecting female mice with 5 IU of pregnant mare serum. 48 h after, females were further injected with 5 IU human chorionic gonadotropin. Immediately after this injection, female mice were mated with male mice overnight; the males were removed the following day. Embryos were collected and genotyped at successive time intervals.

#### Plasmids

pRS416Gal TDP-43 wt-YFP (low copy yeast expression plasmid) was provided by Aaron Gitler, Stanford University (37). The molecular chaperones for the overexpression experiments are produced by the Duennwald Lab and used to transform into pDONR201 (Invitrogen) and further into pAG423Gal-ccdB (high copy yeast expression plasmid) and pAG413Gal-ccdB (low copy yeast expression plasmid) vectors (Addgene plasmid #14149 and 14141) through standard Gateway Cloning protocol. The pEGFP wtTDP-43 plasmid for expression in mammalian cells was provided by Dr. M. Strong, Western University.

#### Antibodies

The antibodies used in this study are shown in Table 1.

**Table 1:** Antibodies used in this study.

| Antigen | Supplier | Use | Concentration |
| --- | --- | --- | --- |
| GFP | Sigma | Western Blot | 1:1000 |
|  | Abcam | Co-IP |  |
| TDP-43 | Sigma | Western Blot | 1:1000 |
|  | Proteintech | Co-IP |  |
| HA | Sigma | Immunofluorescence | 1:1000 |
| Hsp90 | Proteintech | Western Blot | 1:1000 |
|  |  | Immunofluorescence | 1:1000 |
|  | Cell Signaling | Western Blot | 1:1000 |
| Hsp70 | Abcam | Western Blot | 1:1000 |
| Sti1 | In house<br>generated by<br>Bethyl<br>Laboratories<br>Montgomery | Western Blot | 1:5000 |
| Actin | Abcam | Western Blot | 1:1000 |
| PGK-1 | Antibodies-online | Western Blot | 1:2000 |
| Rabbit<br>(Alexa 680) | Life Technologies | Western Blot | 1:2500 |
| Mouse<br>(Alexa 680) | Life Technologies | Western Blot | 1:2500 |
| Rabbit<br>(Alexa 488) | Life Technologies | Immunofluorescence | 1:1000 |
| Mouse<br>(Alexa 488) | Life Technologies | Immunofluorescence | 1:1000 |
| Rabbit<br>(Texas Red) | Life Technologies | Immunofluorescence | 1:1000 |
| Mouse<br>(Texas Red) | Life Technologies | Immunofluorescence | 1:1000 |

## Methods

### Yeast Transformation

Yeast transformations were performed using the standard PEG/lithium acetate method (57). A single colony of yeast is inoculated into 3 mL of yeast-peptone-dextrose (YPD) liquid or selective dextrose (SD) media and incubated at 30°C with shaking overnight. The liquid culture is then combined with 27 mL of YPD liquid to make a 30 mL liquid culture and incubated at 30°C shaking until the cells have reached log phase (an OD_600_ of 0.4 to 0.5). The culture is then centrifuged at 2000 xg for 5 min. The supernatant is aspirated off and discarded and the pellet is washed with 3 mL of sterile water. The cells are centrifuged again at the same speed and time. The pellet is resuspended in 2 mL of 100 mM Li-Acetate in TE buffer after the wash step and incubated at 30°C shaking for 10 min. The culture is centrifuged again after incubation and the pellet is resuspended in 100 μL of Li-Acetate per transformation. Each transformation is composed of 100 μL cell suspension, 250 μL transformation (1 X TE, 40% PEG, and 100mM Li-Acetate), 12 μL salmon sperm DNA, 1 μL (0.3∼0.5 μg) plasmid DNA, and 25 μl DMSO and in the order listed and vortexed thoroughly. The cells then recover at 30°C shaking for 30 min, following a 20-min heat shock at 42°C shaking. After heat shock, the cells are centrifuged for 1 min at 2000 xg, the supernatant aspirated, and the pellet resuspended in 100 μL TE buffer. The cells are then plated onto selective agar plates.

### Yeast Cell Growth Assay

Yeast cell viability is measured using spotting assays. Spotting assays are performed by first inoculating yeast cells in 3 mL in SD media and incubated overnight in a shaking incubator at 30°C. 100 μL of the cells are then taken in an Eppendorf tube to be diluted 1:10 in water to measure the OD_600_, which indicates cell density. In a 96-well plate, we dilute our cell cultures to a cell density normalized to OD_600_ =1 in the first row of wells, followed by a serial dilution of 1:5 in the subsequent 5 wells. We then use a 48-prong Frogger (V&P Scientific) to spot the samples on YPD, SD, and SGal Raf plates lacking selective amino acid markers. The YPD and SD plates are used as growth and spotting controls, whereas the selective galactose raffinose (SGal Raf) plates reflect the toxicity (e.g. of TDP-43) of the induced yeast cells. Incubation periods vary according to the type of plate: YPD plates are incubated 1-2 days; SD plates 2-3 days; and SGal Raf plates 3-5 days. The plates are all incubated at 30°C. The plates are documented during the entire test period to monitor the growth of the yeast colonies by taking photographs using a digital camera.

### Quantification of yeast growth assays

Agar plates were photographed, and converted to black and white images, which were quantified by ImageJ. Unless indicated otherwise, the pixel count for the third dilution of growth assays under the different conditions were used for quantification after subtraction of the background signal. Measured values were input into GraphPad prism was used to generate bar graphs and statistical analyses by applying One Way Analysis of Variance, Tukey Post Hoc using IBM SPSS. Error bars represent standard errors. A minimum of three biological replicas were used for quantification.

### Fluorescent microscopy

Microscopy imaging of YFP tagged constructs was done by first inoculating yeast cells in SD media at 30°C overnight. The cells are then washed twice with sterile water, induced in SGal media, and incubated at 30°C. After induction for 16 to 20 hours, small samples of the culture are placed on a microscope slide. The cells are imaged on Olympus BX-51 Bright Field/Fluorescence Microscope and images were captured using an equipped CCD camera (Spot Pursuit) and the Cytation 5 Cell Imaging Multi-Mode Reader (Biotek).

### Western Blot

A 5 mL yeast culture is first inoculated in SD media overnight. The culture is then centrifuged at 2000 xg, washed twice, and induced in selective galactose (SGal) media overnight. The culture is then centrifuged and washed once with water. The supernatant is discarded and the pellet is resuspended in 200 μL of lysis buffer (50 mM HEPES pH 7.5, 5 mM EDTA, 150 mM NaCl, 1% (v/v) Triton X-100, 50 mM NEM, 2 mM PMSF, and 1X Sigma protease inhibitor tablet) and transferred into an Eppendorf tube. We then add 200 ml of glass beads (ca. 500 mm in diameter) to physically disrupt the cell walls using a vortex mixer for 6 times for 30 sec intervals and cooling on ice for 30 secs between these intervals. We then centrifuge the culture at 5000 xg for 10 min and collect the supernatant in a fresh Eppendorf tube. We perform a BCA Protein Assay to determine the concentration of protein in the sample. The assay was performed according to the Thermo Scientific Pierce BCA Protein Assay Kit instructions. After obtaining the concentration and normalizing the total protein amount in each sample per blot, we dilute the samples with 4x reducing SDS buffer (0.25 M Trisma Base pH 6.8, 8.0% SDS, 40% sterile glycerol, 10% β-mercaptoethanol, 0.04% bromophenol blue).

We run SDS-PAGE with the samples on an 8-16% gradient gel (Bio-Rad Criterion TGX Stain-Free Precast Gels) and/or 12% acrylamide gels at 190 V for about 30 min. The gel is then transferred onto a Nitrocellulose or PVDF membrane (BioRad) using the Bio-Rad Trans-Blot Turbo machine following the manufacturer’s protocol. We then block the membrane using 5% skim milk powder (Carnation) in Phosphate Buffered Saline with 0.01 % (v/v) Tween (PBST) and incubating for 1 h on a shaker. The membrane is then incubated in primary antibody overnight on shaker. Following incubation with the primary antibody, we wash the membrane with 50 ml aliquots of PBST at 10 min intervals for an hour on shaker and then incubate in the secondary antibody for 1 h on a shaker. The membrane is washed again with 50 ml aliquots of PBST in 10 min intervals on shaker for an hour. The membrane is then documented using the ChemiDoc MP System (Bio-Rad) and analyzed using Image Lab (Bio-Rad) and Prism 6 (Graph Pad).

### Sedimentation Assay

The sedimentation assay was adapted from Theodoraki et al. (2012) and Shiber et al. (2013) (51, 52). A 5 mL yeast culture is first inoculated in SD media overnight. The culture is then centrifuged at 2000 xg, washed twice, and induced in SGal media overnight. The culture is then centrifuged and washed once with water. The supernatant is discarded and the pellet is resuspended in 200 μL of lysis buffer (100 mM Tris, pH 7.5, 200 mM NaCl, 1 mM EDTA, 1 mM DTT, 5% glycerol, 0.5% TritonX-100, 50 mM NEM, 2 mM PMSF, and 1X Sigma protease inhibitor tablet) and transferred into an Eppendorf tube. Acid-washed glass beads (425-600 μm, Sigma) are then added to physically lyse the cells using a vortex mixer for 6 times for 30 s intervals and cooling it on ice for 30 s between the intervals. The tubes were pierced with a 16-gauge needle and the lysates (both pellet and supernatant) are collected in a fresh Eppendorf tube by centrifugation in pulses to separate lysates and glass beads. N2a cell lysates were obtained by washing cultured N2a cells in PBS and aspirating off the supernatant and lysing with 100 μL lysis buffer/ 10^6^ cells (150 mM NaCl, 1.0% Triton-X-100, 0.1% SDS, 20 mM Tris pH 7.8-8.0). The cells are then scraped off of the vessel and pipetted into an Eppendorf tube. The total lysates are resuspended and a BCA Protein Assay was performed to determine the concentration of total protein in the sample. The assay was performed according to the Thermo Scientific Pierce BCA Protein Assay Kit instructions. 50 μL of the lysate was mixed with an equal volume of SUMEB Buffer (8 M Urea, 1% SDS, 10 mM MOPS, 10 mM EDTA, and 0.01% bromophenol blue) in a new tube, this aliquot represents total lysates. The rest of the lysate was centrifuged at 500 xg for 15 min at 4°C. 100 μL of the supernatant was transferred into a new tube and mixed with 100 μL of SUMEB buffer, this represents the supernatant portion of the lysate. The remaining supernatant from the lysate was aspirated off. The pellet was resuspended with 100 μL of the lysis buffer (without protease inhibitors) and 100 μL of SUMEB buffer, this represents the pellet portion of the lysate. The samples were boiled at 80°C for 5 min and 25 μL of the samples were loaded onto a 12% acrylamide gel. The gel is then run according to the western blot procedures described above.

### Mammalian Cell Transfection

Mammalian cells were split into dishes or flasks with cell numbers as recommended by manufacturer and incubated for approximately 24 h in DMEM. Cells were grown to approximately 90% confluency and transfected with Lipofectamine LTX with plus reagent (Thermo Fisher) following the supplier recommended concentrations in Opti-MEM (Gibco) low serum media. The cells are incubated at 37 °C for 6 h and followed by a wash in 1X PBS and incubated in DMEM for approximately 20 h at 37 °C. Cells are then split into different dishes or plates for different experimental setups.

### CRISPR-Cas9 Sti1 knock out

Sti1 knockouts of SN56 cells were generated as previously described (54). Sti1 expression was eliminated using CRISPR-Cas 9 (Clustered Regularly Interspaced Short Palindromic Repeats) editing system. The Optimized CRISPR Design Software (http://crispr.mit.edu/) was used to design guide RNAs for the mouse Sti1 gene to transfect SN56 cells (Sti1 Top 1: 5’CACCGGTAGTCTCCTTTCTTGGCGT3’ and Sti1 Bottom 1: 5’AAACACGCCAAGAAAGGAGACTACC3’). The guide RNAs were phosphorylated, annealed and cloned at the BbSI enzyme restriction site into the Addgene vector (px330 modified vector). After this construct was sequenced, Lipofectamine 2000 (Invitrogen) was used to transfect both SN56 cells. Clones were isolated using serial dilution in 96-well plates. Isolated clones were separated until colonies were established. These colonies were then plated in 6-well plates in duplicates. One duplicate was maintained while the other one was lysed for immunoblot analysis. Several clones with decreased Sti1 levels were obtained but only clones with complete elimination of STIP1 expression were expanded and used to investigate the role of Sti1 in TDP-43 mediated toxicity and pathology.

### Mammalian Cell Viability Assay

The CellTiter-Glo Luminescent Cell Viability Assay (Promega) along with the Live/Dead™ Viability/Cytotoxicity Kit (ThermoFisher Scientific, Cat#L3224) were used in this study. The CellTiter-Glo Luminescent Cell Viability Assay delivers a highly sensitive read out of cellular fitness and is particularly useful for studies on the toxicity associated with misfolded proteins. Transfected cells are split into 96 well plates and grown in low glucose and low serum DMEM conditions. This minimal medium increases sensitivity to the toxic effects of protein misfolding in many other systems (71), as well as differentiating N2a cells (69). The cells are incubated at 37°C for 20-24 h. The cell viability assay is then performed according to the supplier’s instructions. Plates were then measured using the Cytation 5 Cell Imaging Multi-Mode Reader (Biotek).

### Immunofluorescence Microscopy

Following transfection, the cells are split into 8 well chambers (Labtek) and grown in low glucose and low serum DMEM at 37 °C for 20-24 h. Each well is seeded with ca. 30,000 cells. The media is then aspirated off and the cells washed 2 times with PBS. 4% paraformaldehyde is then added to the wells to fix the cells for 15 min. This is followed by 3 washes in PBS. The cells are permeabilized with 0.5% TritonX-100 in PBS for 15 min and washed with PBS 3 times, followed by 5 washes in PBS. We block non-specific binding sites with 20% normal goat serum (Gibco) in 0.5% BSA (Sigma) in PBS (PBB) for 45 min. The cells are then washed with PBS 5 times and incubated for 1 h in primary antibody. We wash the cells with PBS 5 times and incubate for an hour in secondary antibody. Following the secondary antibody incubation, we wash in PBS 5 times. All the liquid is then aspirated off and mounted with ProLong® Gold Antifade Mountant with DAPI. The cells are imaged with the Cytation 5 Cell Imaging Multi-Mode Reader (Biotek).

### Co-immunoprecipitation

The N2a cells were lysed according to the lysing protocol described in the sedimentation assay. The lysate is then centrifuged at 15,000 xg for 10 min at 4°C. We discard the pellet and pipet the supernatant into a new Eppendorf tube. We transfer 30 μL of lysate into a new tube, add 30 μL of 2X SDS sample buffer and boil for 5 min. The sample is then frozen. Mouse α-TDP-43 antibody is added to the remaining lysate at a 1:200 ratio, vortexed, and incubated at 4°C overnight on an inverter. We use Bio-Rad SureBeads Protein G Magnetic Beads to perform the pull-down. We transfer 100 μL of beads slurry into a new tube for each lysate sample and wash the beads by resuspending 1000 μL of PBST, magnetizing the beads and discarding the supernatant three times. We add lysate and antibody mixture to the washed beads and incubate lysate-antibody-bead mixture at room temperature on a rocker for 2-3 hours. Following incubation, we wash the pellet-bead mixture three times in PBS and then remove the supernatant. We add 30 μL of 1X SDS sample buffer. We boil our sample for 5 mins and centrifuge bound sample to remove magnetic beads. We also reboil the frozen total lysate. The lysates are loaded onto a polyacrylamide gel. The gel is then run according to the western blot procedures described above. We probe blot with Rabbit α-HA (Sigma) antibody to Co-IP Sti1. We then strip the blot using the Antibody Stripping Buffer (Gene Bio-Application) according to the manufacturer’s protocol and reprobe with α-Hsp90 and α-TDP-43 antibody.

For the SN56 cells, co-immunoprecipitation was performed as described previously (72). Cells were lysed using immunoprecipitation buffer (IP buffer-20 mM Tris, 150 mM NaCl, 1 mM EDTA, 1 mM EGTA, 1% Triton X-100, 0.5% Sodium Deoxycholate, pH 7.5) and incubated in Sepharose beads (abcam, Cat#ab193256) for pre-clearance, and incubated in rabbit polyclonal anti-TDP-43 antibody overnight (Proteintech, Cat#10782-2-AP). In the control condition the SN56 cell lysates were incubated with rabbit polyclonal anti-GFP antibody (Abcam, Cat#ab6556) antibodies after pre-clearance on the orbital shaker overnight at 4°C. The lysates were then incubated in Protein A agarose beads for 4 hours on the orbital shaker at 4°C. The next day, the lysates were centrifuged, the supernatant was discarded. 4x SDS sample buffer (40% glycerol, 240 mM Tris/Hcl pH 6.8, 8% SDS, 0.04% bromophenol blue, 5% β-mercaptoethanol) was added to the beads which were then heated at 95°C for 5 min, centrifuged and then immunoblotted with antibodies against the proteins in question.

### Statistical Analysis

Statistical analysis of viability assays and western blots were done using the GraphPad Prism 6 software. Statistical significance was obtained by performing unpaired t-tests to compare the means and standard deviations between the control data set and the experiment data set (each at a minimum of three biological replicas). Significance levels are indicated using asterisks, where **** is p<0.0001, *** is p<0.001, ** is p<0.01, * is p<0.05. Error bars represent standard errors of the mean.

## Acknowledgements

We would like to thank members of the Prado and Duennwald labs for thoughtful comments on the manuscript, Michael J. Strong for the TDP-43 mammalian plasmid and Jill L. Johnson for providing Sti1 expression vectors.

## Authors ‘contributions

L.T.L., A.R., SED, and BC performed experiments and together with M.L.D., F.B., M.H.L., V.F.P., and M.A.M.P, contributed key reagents, data analyses, interpretation, and critical assessment, L.T.L., A.R., F.B, M.L.D. and M.A.M.P. designed the study and L.T.L.,, M.L.D. and M.A.M.P. wrote the paper. All authors read and approved the final manuscript.

## Availability of data and materials

The data and materials are available from the corresponding author upon request.

## Ethics approval

All procedures for animal use were approved by the institutional Animal Care and Committee at the Western University (2016-103, 2016-104).

## Figure Legends

**S1 Fig: Reduced expression of essential genes Ess1 and Sgt1 do not alter TDP-43 toxicity**. Growth assay of wild type and cells with reduced expression of **A** Ess1 and **B** Sgt1 expressing TDP-43-GFP.

**S2 Fig: High over-expression of Aha1 and Cdc37 is highly toxic in yeast cells. A** Growth assay and **B** fluorescence microscopy of wild type cells highly (2μ) over-expressing Aha1 and Cdc37 with co-expression of TDP-43-GFP. **C** Growth assay of cells lacking Sti1 highly (2μ) over-expressing Aha1 and Cdc37 with co-expression of TDP-43-GFP.

**S3 Fig: Sti1 variants key domains of Sti1 do not alter TDP-43 toxicity**. Growth assays and their quantifications of Sti1 deletion yeast cells co-expressing TDP-43 and variants (amino acid substitutions and truncation) of Sti1.

**S4 Fig: Aha1 and Cdc37 do not alter TDP-43 protein levels in yeast cells**. Immunoblots of **A** high (2μ) over-expression of Aha1 and Aha1 deletion and **B** high (2μ) over-expression of Cdc37 show no change in TDP-43 level when probed with anti-TDP-43 antibody. PGK1 is used as a loading control. The quantitative analysis represents the average of three independent experiments. The vector control samples served as references for the relative changes in the samples with altered Sti1 levels. Error bars represent standard errors of the mean.

**S5 Fig: TDP-43 toxicity and localization are not altered in HEK293 cells. A** Luciferase assay of wild type and CRISPR-Cas9 knock out of Sti1 HEK293 cells expressing TDP-43-GFP do not show change in toxicity. Immunofluorescence of **B** wild type and **C** Sti1 KO cells do not show mislocalization of TDP-43.

**S6 Fig: TDP-43 toxicity and localization are not altered in HeLa cells. A** Luciferase assay of wild type HeLa cells expressing TDP-43-GFP do not show change in toxicity. **B** Immunofluorescence of HeLa cells expressing TDP-43 do not show mislocalization of TDP-43.

